# Effects of Mechanical Loading on Cranial Joint Mesenchymal Stem Cell Proliferation

**DOI:** 10.64898/2026.06.26.734745

**Authors:** Miranda Steacy, Ce Liang, Damith Katudampe Vithanage, Marius Didziokas, Tengyang Qiu, Dale Moulding, Ali Alazmani, Erwin Pauws, Mehran Moazen

**Affiliations:** Department of Mechanical Engineering, University College London, London, UK; Developmental Biology and Cancer Research and Teaching Department, UCL Great Ormond Street Institute of Child Health, University College London, London, UK; School of Mechanical Engineering, University of Leeds, Leeds, UK

**Keywords:** cyclic loading, craniofacial system, mechanobiology, suture, skull

## Abstract

Sutures are the primary sites of cranial bone growth, allowing the skull to accommodate the growing brain. External mechanical stimulus has been shown to delay suture fusion and induce tissue remodelling. Recent studies suggest that *in vivo* cyclic bone loading increased proliferation of mesenchymal stem cells (MSC) in the coronal suture. The overall aim of this study was to understand how many loading sessions (exposure-response) and how long after loading (time-course) did MSC proliferation increase in the coronal suture. In the exposure-response analysis, mice underwent 1, 3, or 5 loading sessions between Postnatal day 7 (P7) and P11, and in the time-course analysis, treated mice underwent 10 loading sessions between P7 and P21. Loading sessions were 10 minutes at a frequency of 1 Hz and a force of 10 g (0.1 N). The loading tip was positioned on the posterior aspect of the left frontal bone, dorsal to the coronal suture. The EdU marker shows a statistically significant increase in proliferation after one loading session and a decrease after three loading sessions. The PCNA marker shows a statistically significant increase after three and five loading sessions. The exposure-response analysis showed that when the results of both markers are combined, levels of proliferation cannot be interpreted until at least five loading sessions have been completed, after which a clear increase in proliferation was observed. In the time-course analysis, proliferation was highest immediately after the final treatment session and 24 hours after the final loading session the effects of mechanical bone loading gradually returned to baseline.

## 1. Introduction

The juvenile craniofacial skeleton is made of several bones joined together via mesenchymal stem cell-filled (MSC) fibrous joints, called sutures. Sutures are the primary sites of cranial bone growth, allowing the skull to accommodate the growing brain and enable skull flexibility during birth (Ang et al., 2022; Flaherty et al., 2016). In addition, sutures can receive and translate external mechanical stimuli directly affecting the overall shape of the skull, suture structure, and suture patency (Herring, 2008; Mao, 2002). Along with morphological changes to the craniofacial complex, an external mechanical stimulus can activate signalling pathways, up- or downregulate gene expression, and alter cellular behaviour (Kopher & Mao, 2003; Peptan et al., 2008; Wang & Mao, 2002).

Several *in vitro* studies have investigated the effects of external forces on craniofacial sutures, finding that tensile forces increase MSC proliferation, maintain suture patency, collagen fibre synthesis, and promote osteoblast differentiation (Ikegame et al., 2001; Tanaka et al., 2000; Tholpady et al., 2007; Yu et al., 2009). However, fewer studies have used *in vivo* models to study mechanobiology of sutures, highlighting that external forces can delay suture fusion, induce craniofacial remodelling, and affect skull growth (Moazen et al., 2022; Peptan et al., 2008; Vij & Mao, 2006). For example, Kopher & Mao (2003) applied cyclic loading to the maxillary incisors of rabbits and found that both tensed nasofrontal and compressed pre-maxillay sutures maintained their patency. We recently showed that minimally invasive *in vivo* cyclic loading of the coronal suture (joining the frontal to the parietal bones) in mouse lead to an increased in MSC proliferation (Steacy et al., 2026). This study’s findings encouraged further investigation into the response time and exposure frequency needed to reach the observed cellular response. Two questions arose regarding the time to effect of this treatment on proliferation. The first being the number of cyclic bone loading sessions required to observe a change in cellular proliferation, an exposure-response analysis. The second is a time-course analysis to determine how long after a full round of treatment (10 loading sessions over 12 days) the cellular proliferation response persists.

The overall aim of this study was to understand how mechanical loading induces proliferation in the coronal suture. Specifically, we hypothesized that cyclic bone loading would gradually increase MSC proliferation with each additional treatment session, and that immediately after a full course of treatment, proliferation would increase, then decrease steadily over the following days.

## 2. Methods

### 2.1. Animals

A total of 35 CD-1 wild type mice were used. In the exposure-response analysis, 18 animals were used with a total of 9 animals in the treated and untreated group. In the time-course analysis, 17 animals were included 8 of which underwent treatment and 9 were used as controls. All the animals were used for histological analyses (n>2)

Note: All animal experiments were approved by the UK Home Office and performed as part of a Project License (number: PP8161503) under the UK Animals (Scientific Procedures) Act 1986. Animal procedures complied with the ARRIVE guidelines and were performed under the supervision of UCL Biological Services.

### 2.2. Mechanical loading

The cyclical loading treatment was adapted from Moazen et al. (2022). An experimental loading set up was developed with an actuator (T-LSR series, Zaber Technologies: res. 50 µm, max. load 200 N) and a force sensor (GSO Series, Transducer Techniques: res. 0.01 N with 1 N capacity) configured to a custom developed LabVIEW program (National Instruments Corp, Austin, TX, USA).

In the exposure-response analysis, animals underwent 1, 3, or 5 loading sessions between postnatal (P) day 7 and P11 and in the time-course analysis, treated mice underwent 10 loading sessions between P7 and P21. Loading sessions were 10 minutes at a frequency of 1 Hz and a force of 10 g (0.1 N). The loading tip was placed on the posterior aspect of the left frontal bone, dorsally to the coronal suture, and lateral to the interfrontal suture (Figure 1).

**Fig 1.**
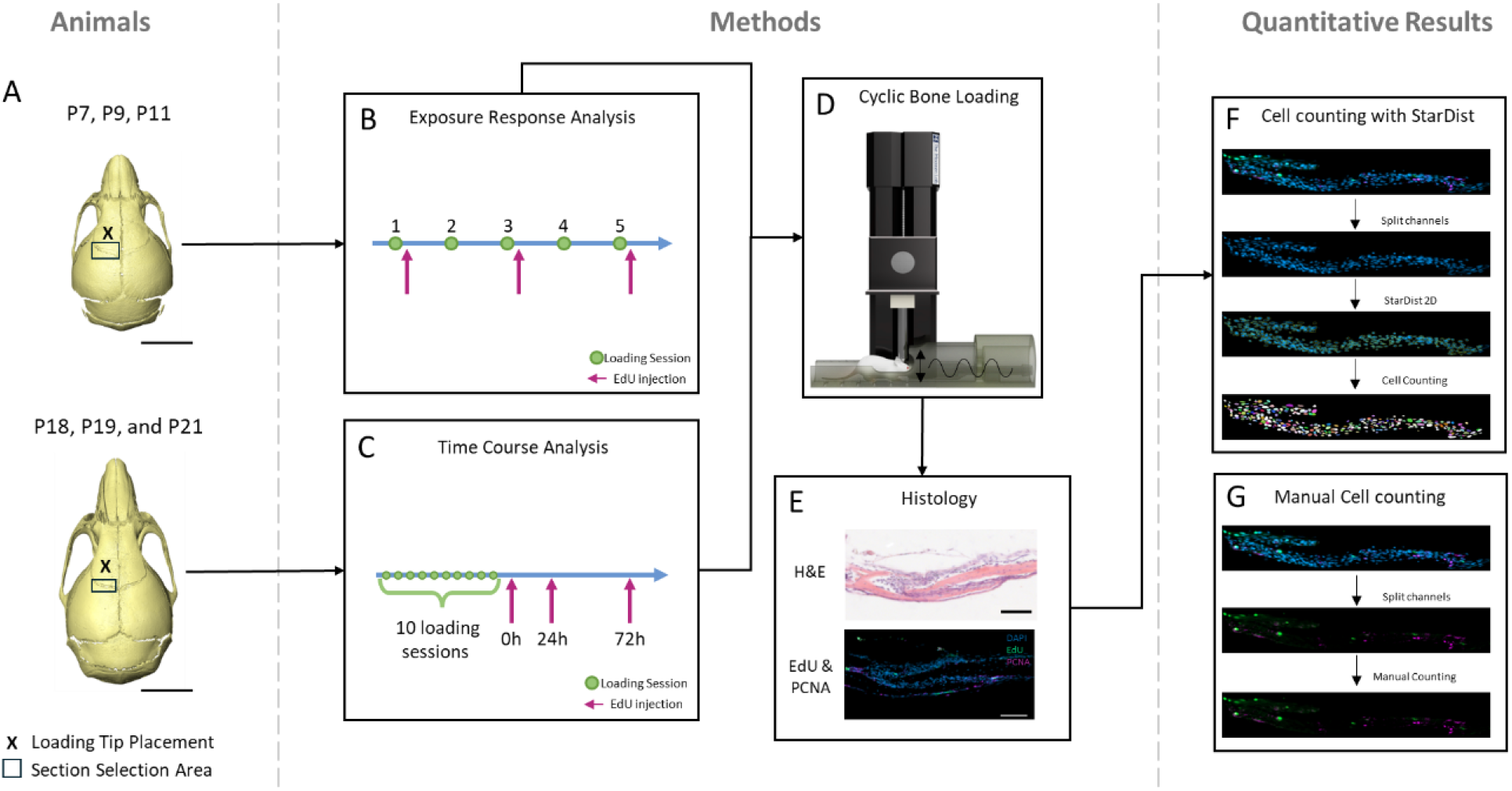
An overview of the experimental workflow. (A) The animals in this experiment included P7, P9, and P11 mice as well as P18, P19, and P21 mice. The loading tip was placed on the posterior aspect of the left frontal bone, lateral to the interfrontal suture and dorsally to the coronal suture. Histology sections included in the proliferation analysis were taken solely from the left coronal suture. (B) The first experiment undertaken was the exposure response analysis. Mice underwent either one (P7), three (P9), or five (P11) loading sessions. Mice were injected with EdU right after their final loading session and collected 4 hours after the IP injection. (C) The second experiment was a time-course analysis. Mice had 10 loading over the course of 12 days and were collected 0 hours after loading (P18), 24 hours after loading (P19), or 72 hours after loading (P21). EdU was injected 4 hours prior to collection. (D) Diagram of the mechanical cyclic bone loading experimental set up. (D) The histology done including H&E and IHC for proliferation markers EdU and PCNA. (F) DAPI stained cell counting of coronal suture cell in ImageJ with StarDist (with CSBDeep). (G) Manual counting of EdU and PCNA-stained cells.

### 2.3. EdU injections

Mice were intraperitoneally injected (50mg/kg body weight) with 5-ethynyl-2′-deoxyuridine (EdU) (baseclick GmbH, Munich, Germany) 4 hours prior to culling.

### 2.4. Histological analysis

Skulls were immediately fixed in 10% formalin at 4°C for 7 days. The fixed skulls were then washed in PBS and decalcified for 10 days in 20% EDTA at 4°C. The skulls were embedded in paraffin and sagittally sectioned 10µm thick. Sections were stained with Haematoxylin and Eosin. Slides were imaged using the Zeiss Axioplan microscope.

### 2.5. Immunohistochemistry

Sagittal sections were then placed in sodium citrate (10mM tri-sodium citrate, 0.5% Tween-20, pH) and underwent antigen retrieval in a steamer. EdU staining was done per the manufacture instructions. Proliferating Cell Nuclear Antigen (PCNA) (clone PC10(3F81), Invitrogen, 1:100) was diluted with PBST (0.01% Tween-20), 10% donkey serum, and 1% bovine serum albumin and applied overnight at 4°C. Secondary antibodies were applied with a 1:300 dilution. DAPI (4,6-diamidino-2-phylindole, Sigma) was pipetted and left for 10 minutes. Immunofluorescence imaging was done on the Zeiss Observer 7 at a 20x 1.0 NA.

### 2.6. Quantitative analysis

DAPI stained images were used to do cell counting in ImageJ (version 1.54i) (n=2 or 3 for each group, 4 technical replicates). An outline of the suture was manually drawn. Cell counting was done by StarDist 2D (with CSBDeep) with a Score Threshold of 0.5 and an overlap threshold of 0.4 (Schmidt et al., 2018). EdU and PCNA-stained cells were manually counted (n>2).

### 2.7. Statistical analysis

For quantitative histological analyses, unpaired t-test with Welch’s correction and Levene’s test were performed. A p-value of <0.05 was considered significant.

## 3. Results

### 3.1. Exposure-response analysis

Mice underwent either one, three, or five loading sessions equating to P7, P9, and P11. After one loading session, within the coronal suture, P7-treated animals had a proportion of proliferative EdU-stained cells of 6.54% (s.d. 1.56%) and PCNA-stained cells of 11.87% (s.d. 1.79%) (Figure 2). Their time-matched control animals had proliferation proportions of 4.83% (1.48%) and 11.60% (s.d. 2.57%) for EdU and PCNA, respectively. The increase in proliferation at P7 after one loading treatment was statistically significant for the EdU proportion (p < 0.01) but not for the PCNA proportion. At P9, after 3 loading sessions, the proportion of EdU-stained cells in the treated animals was 4.16% (1.85%) and PCNA-stained cells was 12.39% (s.d. 2.79%). In the control P9 mice, the proportion of EdU-positive cells in the coronal suture was 6.03% (s.d. 0.84%) and 7.87% (s.d. 2.82%) for PCNA-positive cells. The PCNA-staining at this time point shows a statistically significant increase in proliferation after 3 loading sessions (p < 0.001). In contrast, the EdU-staining shows a statistically significant decrease in proliferation after 3 loading sessions (p < 0.008). Finally, at P11 and after 5 loading sessions, treated animals had proliferation proportions of 6.38% (s.d. 0.84%) and 13.82% (s.d. 3.72%) for EdU and PCNA, respectively. The control P11 animals had proliferation proportions of 5.60% (s.d. 2.16%) and 9.43% (s.d. 2.39%). The increase in PCNA-positive cells in the treated animals was statistically significant after five loading sessions (p < 0.002).

**Fig 2:**
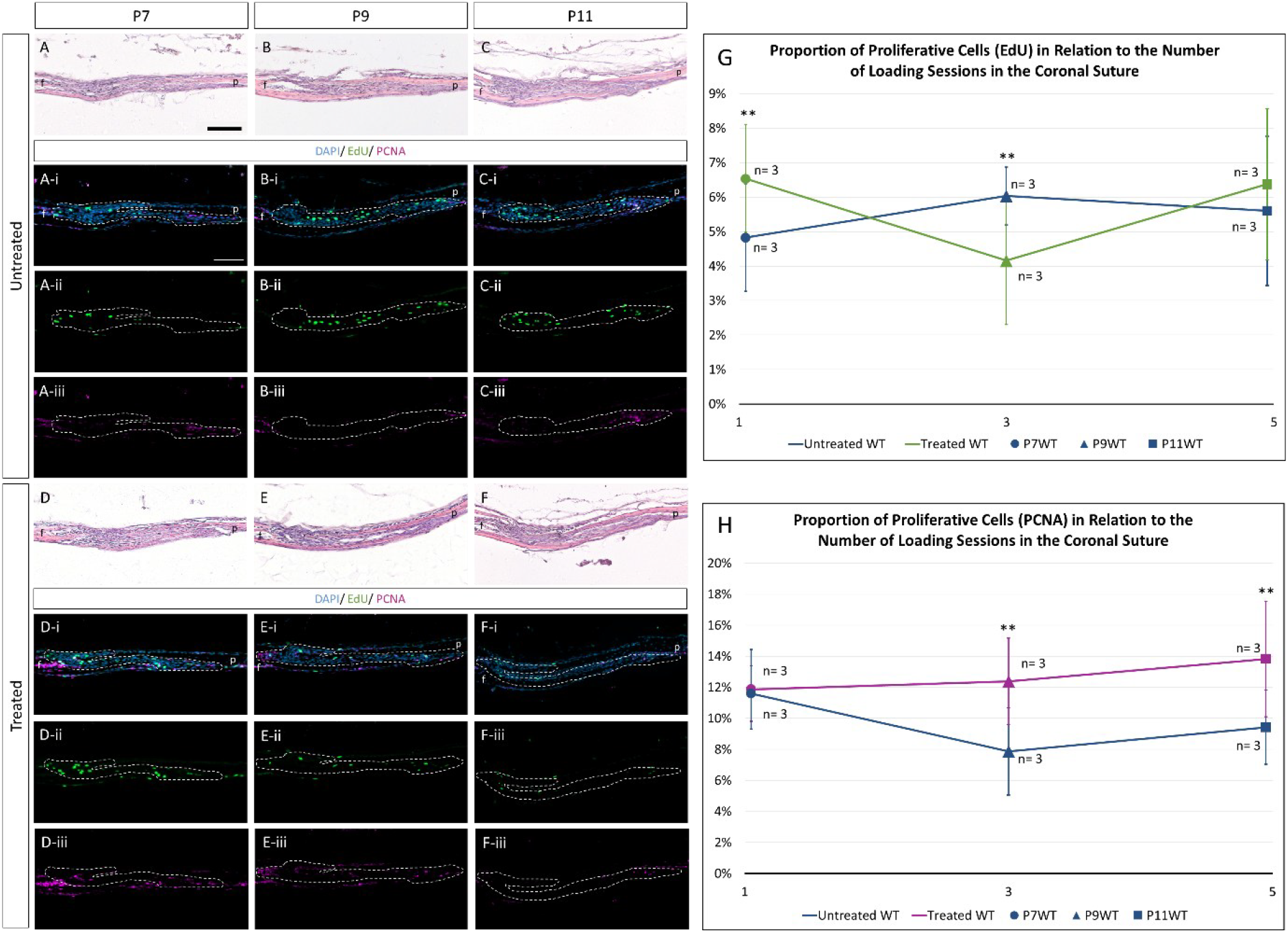
Exposure-response analysis at P7 (one loading session), P9 (three loading sessions), and P11 (five loading sessions). (A-C, D-F) H&E-stained sagittal sections for the coronal suture in treated (loaded) and control animals. (i) Immunohistochemistry composites of sagittal sections for the coronal suture stained for DAPI (blue), EdU (green), and PCNA (magenta). Dashed line outlining the coronal suture. (ii) Single channel immunohistochemistry of EdU. (iii) Single channel immunohistochemistry of PCNA. f: frontal, p: parietal. Scale bar: 100 μm. (G) Exposure-response graph showing the proportion of proliferative cells (EdU) in the coronal suture cells after one, three, and five loading sessions. (H) Exposure-response graph showing the proportion of proliferative cells (PCNA) in the coronal suture cells after one, three, and five loading sessions. (n=3, 4 technical replicates per group) (** p < 0.01)

### 3.2. Time-course analysis

After two weeks of cyclic mechanical bone loading, mice were collected immediately after treatment (0 hours, P18), one day after treatment (24 hours, P19), or three days after treatment (72 hours, P21) (Figure 3). P18 mice had an average EdU proliferation percentage of 3.41% (s.d. 2.18%), whereas their age-matched controls had 1.24% (s.d. 0.89%) EdU-positive cells in the coronal suture. The P18-treated animals had an average of 22.17% (s.d. 3.87%) positive PCNA-stained cells. The age-matched controls had a proportion of PCNA-positive cells of 15.51% (s.d. 5.32%). At P18, the increase in EdU and PCNA-positive cells in the treatment group was statistically significant (p <0.02 and p < 0.008, respectively). In the P19 mice, treated animals had an EdU-positive cell percentage of 2.93% (s.d. 2.17%) and a PCNA-positive cell percentage of 23.85% (s.d. 6.22%). P19 untreated mice had an EdU staining percentage of 2.11% (s.d. 1.13%) and a PCNA staining percentage of 16.57% (s.d. 4.69%). In P19 mice culled 24 hours after the last loading session, the increase in proliferation was statistically significant for the PCNA marker (p < 0.01), whereas the increase in EdU-positive cells was not statistically significant. Mice that were culled 72 hours after treatment at P21 had an EdU proliferation percentage of 2.80% (s.d. 1.01%), and in untreated animals, 1.84% (s.d. 0.58%). The P21-treated animals had a PCNA-positive percentage of 16.30% (s.d. 4.13%), and untreated animals had a PCNA-positive percentage of 15.10% (s.d. 2.82%). Proliferation was not statistically significant at P21, 72 hours after treatment.

**Fig 3:**
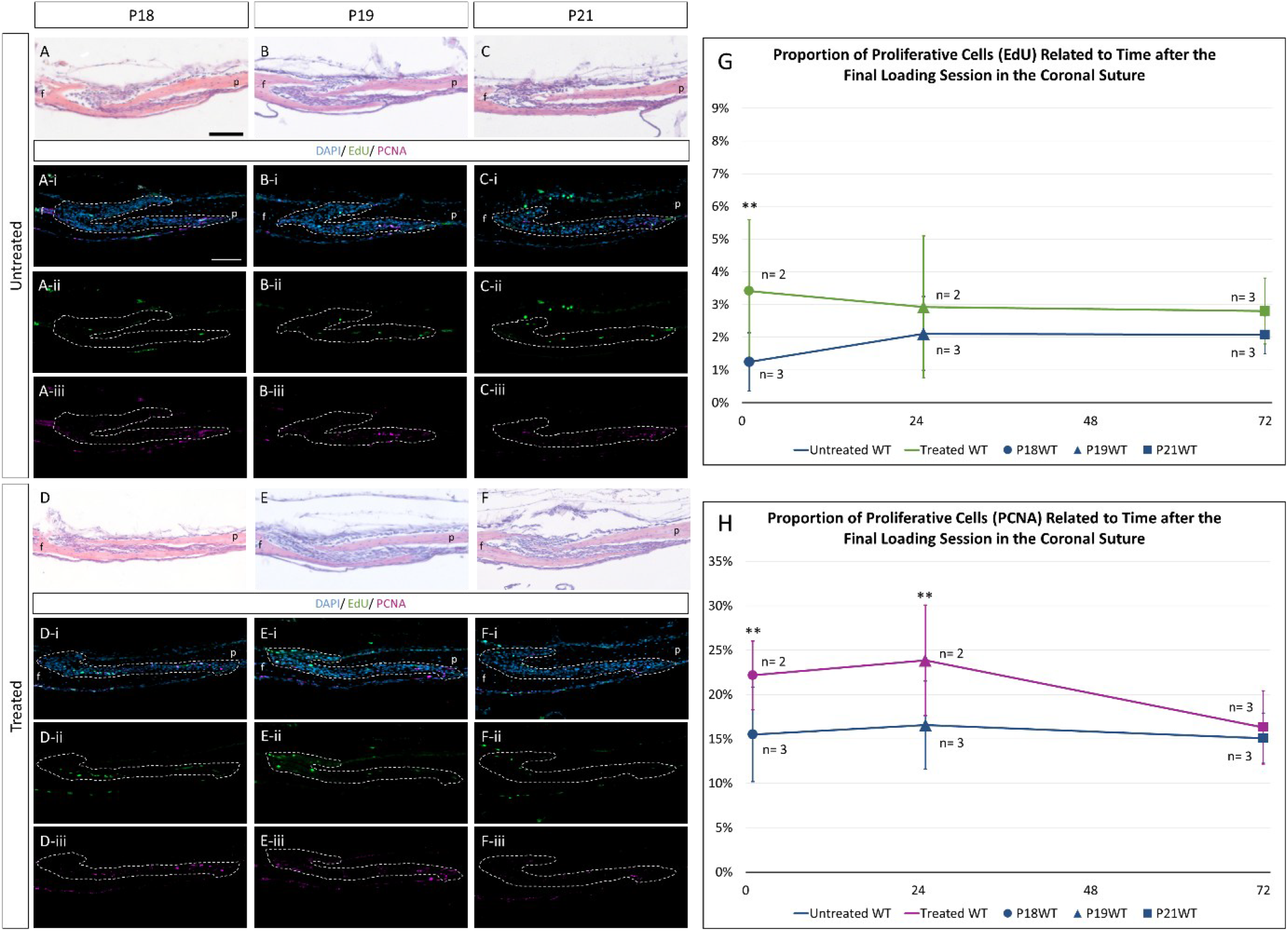
Time-course analysis at P18 (0 hours after 10 loading sessions), P19 (24 hours after 10 loading sessions), and P21 (72 hours after 10 loading sessions). (A-C, C-F) H&E-stained sagittal sections for the coronal suture in treated (loaded) and control animals. (i) Immunohistochemistry composites of sagittal sections for the coronal suture stained for DAPI (blue), EdU (green), and PCNA (magenta). Dashed line outlining the coronal suture. (ii) Single channel immunohistochemistry of EdU. (iii) Single channel immunohistochemistry of PCNA. f: frontal, p: parietal. Scale bar: 100 μm. (G) Time-course graph showing the proportion of proliferative cells (EdU) in the coronal suture cells 0 hours, 24 hours, and 72 hours after 10 loading sessions. (H) Time-course graph showing the proportion of proliferative cells (PCNA) in the coronal suture cells 0 hours, 24 hours, and 72 hours after 10 loading sessions. (n=2 or 3, 4 technical replicates per group) (** p < 0.01)

## 4. Discussion

Our recent study found that cyclic loading of coronal suture increases MSC proliferation in the coronal suture (Steacy et al., 2026). This study aimed to further investigate the exposure frequency and response time needed to reach peak MSC proliferation. The main finding of the exposure-response analysis was that cyclic bone loading increased EdU-positive cells after one loading sessions and decreased after three loading sessions, whereas PCNA-positive cells increased after three and five loading sessions. In the time-course analysis, the main finding from both the EdU and PCNA markers was that immediately after the final treatment session, proliferation peaked, and 24 hours after the final loading session, the effects of mechanical bone loading gradually returned to baseline.

EdU labels newly synthesized DNA during the S-phase of cell division, whereas PCNA is abundant in cells during the G1, S, G2, and M phases (Mead & Lefebvre, 2014). The higher percentage of PCNA-positive cells seen in the coronal suture can be explained by its presence throughout a greater proportion of the cell cycle, its persistence half-life of around 20 hours, meaning it remains in the cell for 24 to 48 hours in newly quiescent cell, and its transient accumulating at sites of DNA damage (Chapman & Wolgemuth, 1994; Essers et al., 2005; McCarty, 2007).

The main discrepancy between the EdU and PCNA markers in this study lies in the exposure-response analysis. After 3 loading sessions, EdU shows a decrease in proliferation, whereas PCNA shows an increase. The EdU result was inconsistent with the initial hypothesis that cyclic bone loading would gradually increase MSC proliferation with each additional treatment session. Both markers are widely used and considered highly reliable (Mead & Lefebvre, 2014). Therefore, neither proliferation marker can be discounted solely on the results from this study. One explanation for this difference could be the low number of animals included in this study. Kulldorff et al., (2000) found that the correlations between BrdU and PCNA assays vary substantially based on the number of biopsies obtained for each individual, emphasising the importance of having a large number of participants in cell proliferation measurement studies. Altogether, the exposure-response analysis showed that when the results of both markers are combined, levels of proliferation cannot be interpreted until at least five loading sessions have been completed, at which point a significant increase in PCNA and a non-significant increase in EdU proliferation is observed. Tholpady et al., (2007), found similar results, when sutures were artificially maintained open by tensile force the increase in proliferative cells was not apparent as early as 1 week but was very clear in the second week.

PCNA showed that loading had a cumulative effect on proliferation. Proliferation was 11.87%, 12.39%, and 13.82% after one, three, and five loading sessions, respectively. Kaspar et al., (2002) found that in human bone-derived osteoblast-like cells, proliferation increases with the number of applied cycles until a maximum cycle number is reached. Exposure-response analyses tend to emphasize the ‘therapeutic window’, highlighting the cumulative, plateauing then adverse effects of external loading. Guo, (2026) illustrates this with a step line showing the gradual increase in proliferation, a window of 3 to 14 days, and a plateau thereafter. In this study, the treatment remained in the ‘therapeutic window’ and the time-course analysis corroborated the sustained positive effects of the treatment protocol.

The time-course analysis, where proliferation was sustained for 24 hours after the last treatment session, aligns with the hypothesis and other studies investigating the onset and duration of proliferation after loading. In explant culture and *in vivo*, cell proliferation increases upon tensile strain for 24 hours and declines 72 hours post-treatment (Hickory & Nanda, 1987; Mizukoshi et al., 2021).

The main limitation of this study was the number of animals included. Due to the labour-intensive nature of the cyclic bone loading, only nine animals could be treated in one day. In the future, it would be important to repeat this protocol on more animals to increase the n-numbers, which may shed light on the EdU and PCNA marker discrepancies observed in the exposure-response analysis.

In conclusion, cyclic bone loading of the coronal suture showed an increase in proliferation after a minimum of five loading sessions and had a cumulative effect with each loading session. Cyclic bone loading also showed the highest level of proliferation in the coronal suture right after the final loading session in a ten session loading treatment protocol which decline 24 hours after the last session.

## 5. Declaration

All authors contributed to the study’s conception and design. Specimen preparation was performed by Miranda Steacy, Ce Liang, Damith Katudampe Vithanage, Marius Didziokas, Tengyang Qiu, and Erwin Pauws. The custom loading set up was developed by Ali Alazmani. All histology and image processing was performed by Miranda Steacy with the guidance of Dale Moulding. All statistical analysis was done by Miranda Steacy. The first draft of the manuscript was written by Miranda Steacy. All authors commented on previous versions of the manuscript. All authors read and approved the final manuscript. Mehran Moazen and Erwin Pauws supervised the project and secured the funding.

## 6. Funding

This work was supported by the Engineering and Physical Science Research Council (EP/W008092/1; EP/R513143/1 – 2592407 and EP/T517793/1 - 2592407) and the NIHR Great Ormond Street Hospital Biomedical Research Centre.

## Notes

### Competing Interest Statement

The authors have declared no competing interest.

